# Purification and biochemical characterization of the DNA binding domain of the nitrogenase transcriptional activator NifA from *Gluconacetobacter diazotrophicus*

**DOI:** 10.1101/2023.05.30.542961

**Authors:** Heidi G. Standke, Lois Kim, Cedric P. Owens

**Author notes:** **Corresponding Author** Cedric Owens, Chapman University, Schmid College of Science and Technology, Keck Center for Science and Engineering 226, One University Drive, Orange, CA 92866.

## Abstract

NifA is a σ^54^ activator that turns on bacterial nitrogen fixation under reducing conditions and when fixed cellular nitrogen levels are low. The redox sensing mechanism in α-proteobacterial NifA is poorly understood. In this work, we examine if a Cys pair that is part of a C(X)_5_C motif and located immediately upstream of NifA’s DNA binding domain is involved in redox sensing in NifA from the α-proteobacterium *Gluconacetobacter diazotrophicus* (*Gd*). We hypothesize that the Cys residues’ redox state may directly influence the DNA binding domain’s DNA binding affinity and/or alter the protein’s oligomeric sate. Two DNA binding domain constructs were generated, a longer construct (2C-DBD), consisting of the DNA binding domain with the upstream Cys pair, and a shorter construct (NC-DBD) that lacks the Cys pair. The *K*_d_ of NC-DBD for its cognate DNA sequence (nifH-UAS) is equal to 20.0 μM. The *K*_d_ of 2C-DBD for nifH-UAS when the Cys pair is oxidized is 34.5 μM. Reduction of the disulfide bond does not change the DNA binding affinity. Additional experiments indicate that the redox state of the Cys residues does not influence the secondary structure or oligomerization state of the NifA DNA binding domain. Together, these results demonstrate that the Cys pair upstream of the DNA binding domain of *Gd*-NifA does not regulate DNA binding or domain dimerization in a redox dependent manner. This suggests that other Cys residues in NifA, such as those located in the central AAA^+^ domain, are responsible for redox sensing.

## Introduction

In proteobacteria, NifA is the central regulator of bacterial nitrogen fixation, the conversion of dinitrogen gas (N_2_) into ammonia (NH_3_). NifA regulates the expression of the nitrogenase structural genes *nifH, nifD*, and *nifK* as well as other enzymes that support N_2_ reduction [1-4]. Like most σ^54^ activators, NifA has a three-domain architecture (Figure 1A), consisting of an N-terminal receiver domain, a central AAA^+^ domain, and a C-terminal DNA binding domain [5-7]. The DNA binding domain consists of a tri-helical helix-turn-helix (HTH) domain [8] and is connected to the AAA^+^ domain through a flexible interdomain linker (IDL).

**Figure 1.**
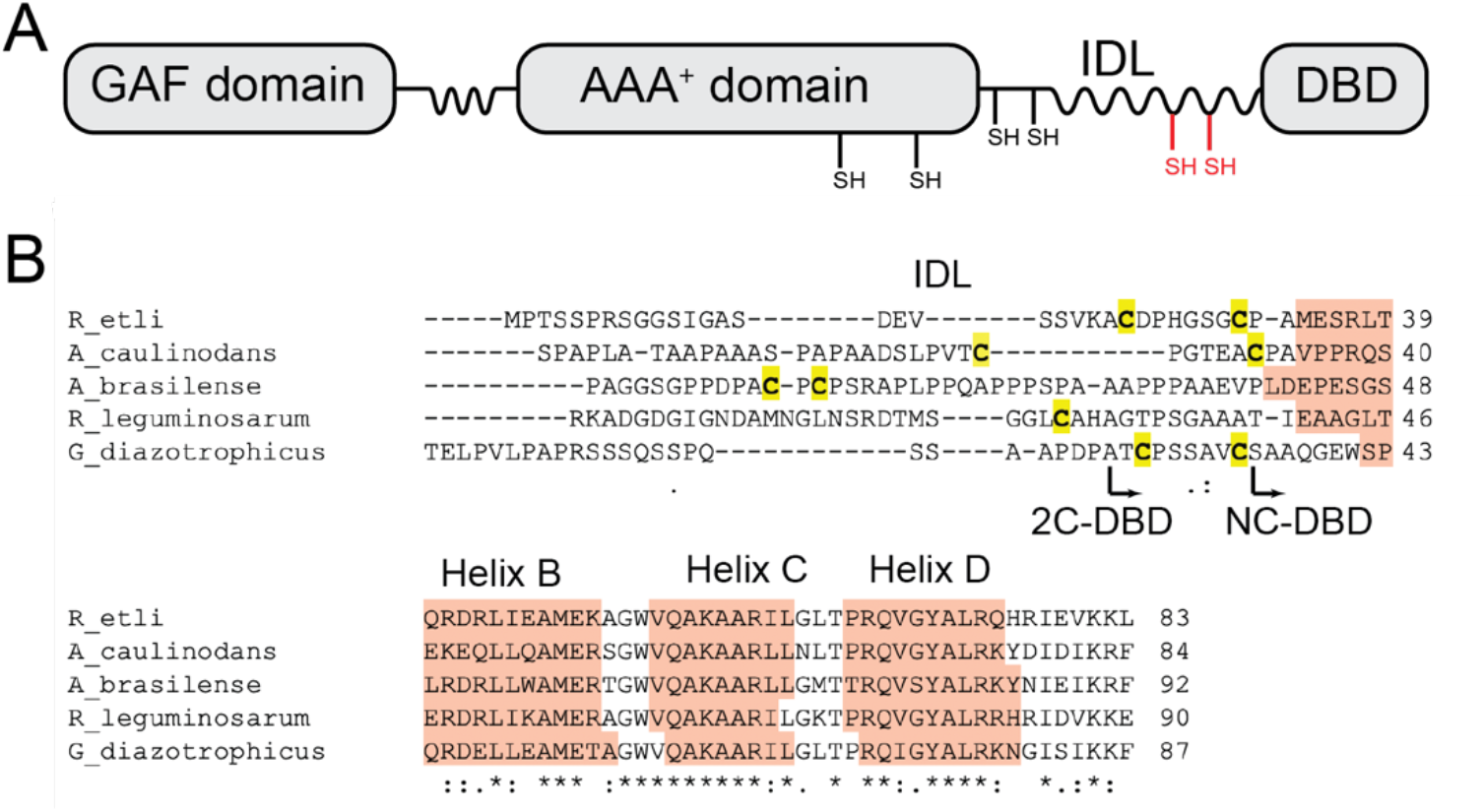
(A) Domain architecture of NifA. In proteobacteria, the NifA receiver domain is a GAF domain. Conserved Cys residues in the AAA^+^ domain and start of the IDL are colored in black. Regions of the protein that are predicted to be flexible are depicted as wavy lines. The location of the Cys pair immediately upstream of the DNA binding domain (DBD) in *G. diazotrophicus* NifA is shown in red. (B) Representative sequence alignment of several NifA IDLs and DNA binding domains from α-proteobacteria highlighting the location of Cys residues upstream of the tri-helical HTH domain. Even though the overall sequence in the IDL is not conserved, the presence of Cys residues is widespread. Helices are named based on the nomenclature proposed by Vidangos et. al. [8] in which NifA-like proteins lack “Helix A”. The boundaries for the *G. diazotrophicus* NC-DBD and 2C-DBD constructs are indicated by arrows.

Activation of nitrogen fixation genes by NifA occurs when fixed cellular nitrogen levels are low and intracellular redox levels are reducing [1]. To activate transcription, NifA undergoes a conformational change from a dimer to a hexamer. The hexamer interacts with the RNA polymerase σ^54^ factor RpoN and initiates transcription in an ATP-dependent manner [6, 9]. Nitrogen sensing in NifA occurs in the receiver domain [10, 11]. There are two distinct mechanisms of NifA redox sensing depending on the source organism. In γ-proteobacteria, redox sensing takes place on a separate protein, NifL. When oxygen levels are high, NifL interacts with NifA, inhibiting ATP hydrolysis in the AAA^+^ domain which prevents transcriptional activation [1, 12]. In contrast, in α- and β-proteobacteria, redox sensing takes place on NifA itself [13, 14]. NifA contains four conserved cysteine residues in the AAA^+^ domain and at the start of the IDL (Figure 1A) [14, 15]. In addition, NifA in many α-proteobacterial species contain a single Cys or two Cys residues upstream of the DNA binding domain (Figure 1C) at the C-terminal end of the IDL. Together, this suggests that NifA may sense redox conditions using a metal cluster or though reversible disulfide bond formation. However, to date, the mechanism for NifL-independent NifA redox sensing is unknown, especially how redox changes at the Cys sites relate to DNA binding, NifA oligomerization, or ATP hydrolysis.

The presence of Cys residues immediately upstream of the DNA binding domain in α- and β-proteobacteria (Figure 1B) is intriguing. DNA binding domains of σ^54^ activators bind to their palindromic target sequences as dimers [8, 16]. The Cys residues are located approximately along the predicted dimerization interface, suggesting that the redox state of the thiols may influence the DNA binding domain’s structure to enable stronger DNA binding under reducing conditions. Such an environmental sensing mechanism, occurring directly in the DNA binding domain itself, would represent a novel mechanism for σ^54^ activators.

To determine the role of the Cys residues, we recombinantly expressed and purified the DNA binding domain of NifA from the α-proteobacterium *Gluconacetobacter diazotrophicus* (*Gd*), which has a well-characterized nitrogen fixing system [17-19]. The DNA binding domain of *Gd*-NifA is homologous to that of *Herbaspirillum seropedicae* NifA, which was shown to bind DNA independently of the rest of the protein [20]. *Gd*-NifA contains a pair of Cys residues preceding the DNA binding domain (Figure 1C), indicating that the domain could form either inter or intramolecular disulfide bonds. We generated two DNA binding constructs, 2C-DBD, which is composed of the DNA binding domain and the part of the IDL containing the Cys pair, and NC-DBD, which only contains the core DNA binding domain (Figure 1C). Biophysical characterization of 2C-DBD and NC-DBD indicates they binds to DNA with similar affinity, however, there was no evidence that the Cys residues mediate dimerization or have a significant role in altering the DNA binding domain structure and DNA binding affinity in a redox-dependent manner.

## Materials and Methods

### Reagents

Reagents were purchased from Sigma Aldrich, Fisher Scientific and were ACS grade or equivalent. Cloning reagents were purchased from New England Biolabs or Fisher Scientific.

### Molecular cloning of NC-DBD and 2C-DBD

NC-DBD and 2C-DBD were amplified from a previously generated plasmid containing full-length NifA (*G. diazotrophicus* NifA-pMAL-c5x, Owens laboratory, unpublished results). The forward primers for were 5’-CGC GCT AGC TCG GCC GCG CAG GGG and 5’-CGC GCT AGC GCG ACG TGC CCG for NC-DBD and 2C-DBD, respectively. The reverse primer for both constructs was 5’-CGC GGA TCC TCA GAA TTT CTT GAT GGA AAT CCC. The forward primers contain an NheI restriction enzyme recognition site, whereas the reverse primer contains a BamHI site. PCR was performed with a denaturation temperature of 95 °C, an annealing temperature of 67 °C, and an extension temperature of 72 °C for 30 cycles. The amplified PCR product was then purified on a 1% agarose gel and extracted using a Thermo Scientific GeneJet PCR purification kit. The purified PCR product and pET28a plasmid were incubated with NheI and BamHI restriction enzymes (NEB) following the manufacturer’s protocol for 3 hours. After restriction digest, shrimp alkaline phosphatase (Affymetrix) was added for 30 minutes to pET28a. Digested PCR product and pET28-a plasmids were then run on a 1% gel and purified using Thermo Fisher Scientific’s GeneJet PCR purification kit. NC-DBD and DBD 2C-DBD were ligated into pET28-a using T7 ligase (NEB) and subsequently transformed into chemically competent *E. coli* 5α cells (NEB) via heat shock and plated on LB medium containing kanamycin at a concentration of 50 μg/mL. Several colonies were then transferred into 5 mL of liquid LB culture containing 50 μg/mL kanamycin and grown overnight. The respective plasmids were then purified using a Thermo Fisher Scientific GeneJET Plasmid Miniprep Kit and sent for sequencing (Genscript).

### Protein expression

NC-DBD and 2C-DBD were transformed into *E. coli* BL21 using standard heat shock protocols. A single colony was selected and grown overnight at 37 °C and 250 rpm in 100 mL LB broth containing 50 μg/mL kanamycin. The next day, 25 mL of overnight culture was added per L of LB broth containing 30 μg/mL kanamycin, and the cells grown at 37 °C and 250 rpm. Expression was induced by addition of IPTG to a final concentration of 0.4 mM when the optical density reached 0.6 - 0.9. Expression was allowed to occur for four hours after which the cells were spun down at 5000 rpm. Cell pellets were stored at -20 °C until use.

### Purification of NC-DBD and 2C-DBD

Cells were resuspended in a wash buffer containing 50 mM Tris, pH 8, 500 mM NaCl, 20 mM imidazole, 1 mM PMSF, 10 mM BME, and a pinch of lysozyme. Cells were lysed by sonication in an ice bath (four cycles of 30 seconds with 30 second breaks between cycles) and the cell free extract spun down at 12,500 rpm. The supernatant was loaded onto a HiLoad Ni^2+^ column (GE healthcare) and the protein eluted using a linear gradient with 50 mM Tris, pH 8, 500 mM NaCl, 500 mM Imidazole. Protein purity was verified by 15% SDS-PAGE and fractions containing the DNA binding domain were pooled. The protein was then extensively dialyzed against 10 mM Tris, pH 8, and 60 mM NaCl. If necessary, the protein was further purified on an S75 10/300 gel filtration column (GE healthcare) equilibrated with 10 mM Tris, pH 8, and 60 mM NaCl.

The His-tag was removed via thrombin cleavage using Biovision Thrombin-agarose beads, where the protein concentration was 1 mg/mL during cleavage. The His-tag was separated from DBD on an S75 10/300 GE gel filtration column (GE healthcare) equilibrated with 10 mM Tris, pH 8, and 60 mM NaCl. SDS-PAGE was run to confirm His-tag cleavage and protein purity and identify fractions containing 2C-DBD and NC-DBD. The protein was subsequently pooled, concentrated and stored at -80 °C until use. Protein concentration was determined using ε_280nm_ equal to 12,490 M^-1^cm^-1^ for NC-DBD and 12,553 M^-1^cm^-1^ for 2C-DBD.

### MALDI-TOF of NC-DBD and 2C-DBD

NC-DBD and 2C-DBD at concentrations between 70-200 μM were diluted 10-fold in a 1:10 mixture of sinapinic acid and water and performed using methods described in reference [21].

### Analytical gel filtration

2C-DBD and NC-DBD samples were run on a S75 10/300 column equilibrated with 10 mM Tris, pH 8 and 60 mM NaCl. The flow rate was 0.7 mL/min. The protein concentration was typically 5 mg/mL, but lowered up to 0.5 mg/mL to test the concentration dependence of the elution volume. To achieve reducing conditions, TCEP (5 mM) or DTT (10 mM) was added to the buffer, and the protein was incubated for 10 min prior to being run on a S75 10/300 column equilibrated with 10 mM Tris, pH 8 and 60 mM NaCl, and 5 mM TCEP or 10 mM DTT.

### Glutaraldehyde crosslinking

Crosslinking was carried out in 25 mM HEPES, pH 8, 25 mM NaCl. The protein concentration was 0.2 mg/mL and the final glutaraldehyde concentrations were 0.1% or 0.01%, as indicated in the figure. Crosslinking proceeded for 5 min and was quenched by addition to Tris, pH 8 to a final concentration of 200 mM. Samples were denatured and resolved by 15% SDS-PAGE.

### Circular dichroism spectroscopy

Circular dichroism (CD) spectroscopy was performed on 2C-DBD and NC-DBD in 1 mM Tris, pH 8, 6 mM NaCl. Scans were taken at the temperatures indicated in the main text with an integration time of 2 sec, a bandwidth of 1 nm, a data pitch of 0.2 nm, and a scanning speed of 100 nm/min. Each spectrum consists of the average of four acquisitions. For thermal denaturation experiments, ellipticity was monitoring at 222 nm as the temperature was increased linearly with a ramp rate of 5 °C/min between 4 °C to 90 °C. In thermal unfolding experiments, data was converted into percent unfolded using following formula: Percent unfolded = (θ – θ_4°C_)/(θ_94°C_ – θ_4°C_) ×100%.

### Fluorescence anisotropy measurements

The DNA probe (IDT) consisted of 900 nM nifH-UAS duplex with six flanking nucleotides on each side (5’ - CGG TTT TGT CAG GCT TCG CAC AAA GCC G - 3’) that was fluorescently labeled with a TAMRA fluorophore at the 5’ end of the forward strand. DBD constructs were added to the DNA probe at concentrations between 0-80 μM. The DNA binding buffer contained 10 mM Tris, pH 8, 60 mM NaCl, and 0.2 mM MgCl_2_. When reducing conditions were desired, DTT was added to a final concentration of 5 mM. Control experiments indicated that DTT, at 5 mM concentration, does not alter the fluorescent properties of the TAMRA probe, and, furthermore, that NC-DBD and 2C-DBD do not quench probe fluorescence. DNA and DNA binding proteins were incubated for 30 minutes at room temperature and then transferred to a 384 well plate (Corning). Fluorescence anisotropy was measured using an excitation wavelength of 500 nm and an emission wavelength of 577 nm on a Tecan Spark plate reader fitted with a 50% dichroic 510 mirror. Anisotropy-based binding curves was fit in Graphpad Prism to a One Site Binding Curve equation, *r* = *r*_o_ + *B*_max_/(*K*_D_ + [DNA]), where *r* = is the measured anisotropy value, *r*_o_ the initial anisotropy of the probe by itself, *B*_max_ is the maximum specific binding, and *K*_d_ is the binding constant.

## Results

### Structural analysis of NifA DNA binding domain models

Structural modeling was used to predict the structure of *Gd*-NifA’s DNA binding domain. Modeling was carried out using Robetta [22]. First, we predicted the structure of NC-DBD (Figure 2A), which comprises only the DNA binding domain without the IDL (*Gd*-NifA residues 530 to 581). NC-DBD features a tri-helical HTH motif (Helices B, C, and D according to the nomenclature proposed by Vidangos et. al. [8]) and is homologous to the DNA binding domain of NtrC1 with an RMSD of 0.8 Å (Figure 2B). Next, a structural prediction of full length NifA was generated to obtain insights into the boundaries and possible structure of the IDL. The IDL does not feature a defined secondary structure and is predicted with low accuracy, indicating that NifA does not feature “Helix A” upstream of the HTH motif. This makes NifA similar to NtrC1 but unlike NtrC and NtrC4 which both feature Helix A [8]. The structural model of 2C-DBD, comprising residues 520-581 of *Gd*-NifA, is shown in Figure 2C. The Cys pair is modeled as being oxidized. In the 2C-DBD model, the IDL is aligned approximately colinearly to Helix B. Such an orientation would point the Cys residues away from a neighboring protomer in a DNA-bound NifA dimer (Figure S1). However, since the IDL is flexible, it can sample other orientations. We investigated possible conformation of the IDL using the MoMA loop modeling server [23], which indicated that the Cys containing IDL region can access conformations that would allow it to form an intermolecular disulfide. Figure S1 shows a possible 2C-DBD dimer structure in which Cys residues from the two protomers are in proximity.

**Figure 2.**
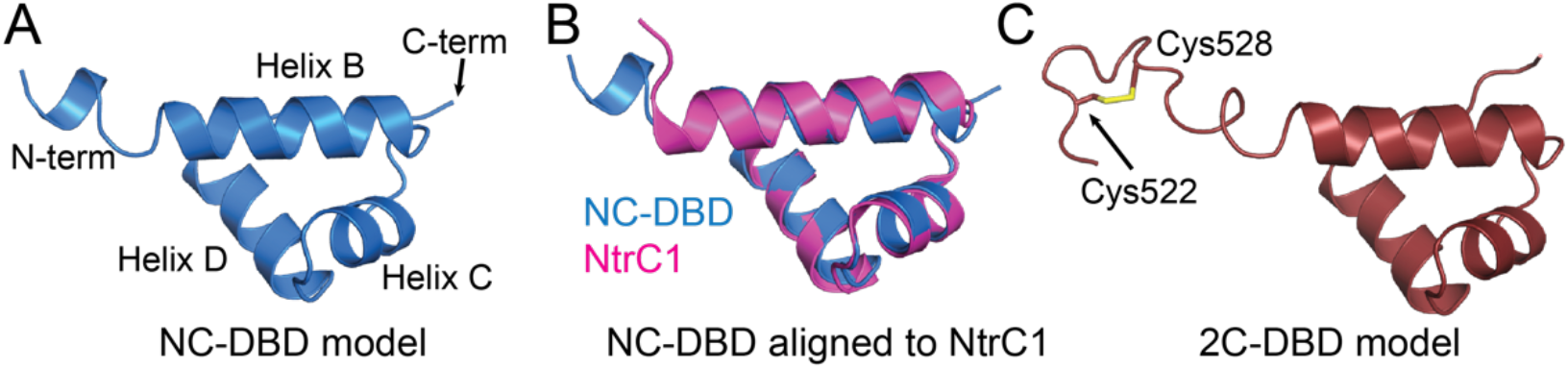
(A) Structural model of NC-DBD. (B) Structural alignment of NC-DBD and the DNA binding domain of NtrC1 (pdb id: 4l5e), and (C) model of 2C-DBD.

### Protein purification and oligomeric state analysis

NC-DBD and 2C-DBD were expressed as His-tagged fusion proteins in *E. coli* BL21 cells and purified in two steps by Ni^2+^ affinity and gel filtration chromatography. The His-tag was cleaved with thrombin and separated from the DNA binding domain by gel filtration chromatography. SDS-PAGE analysis indicates that after purification both 2C-DBD and NC-DBD were homogeneous, and that His-tag cleavage was complete (Figure 3A). The molecular weight of cleaved NC-DBD and 2C-DBD was further confirmed by MALDI-TOF (Figure S2).

**Figure 3.**
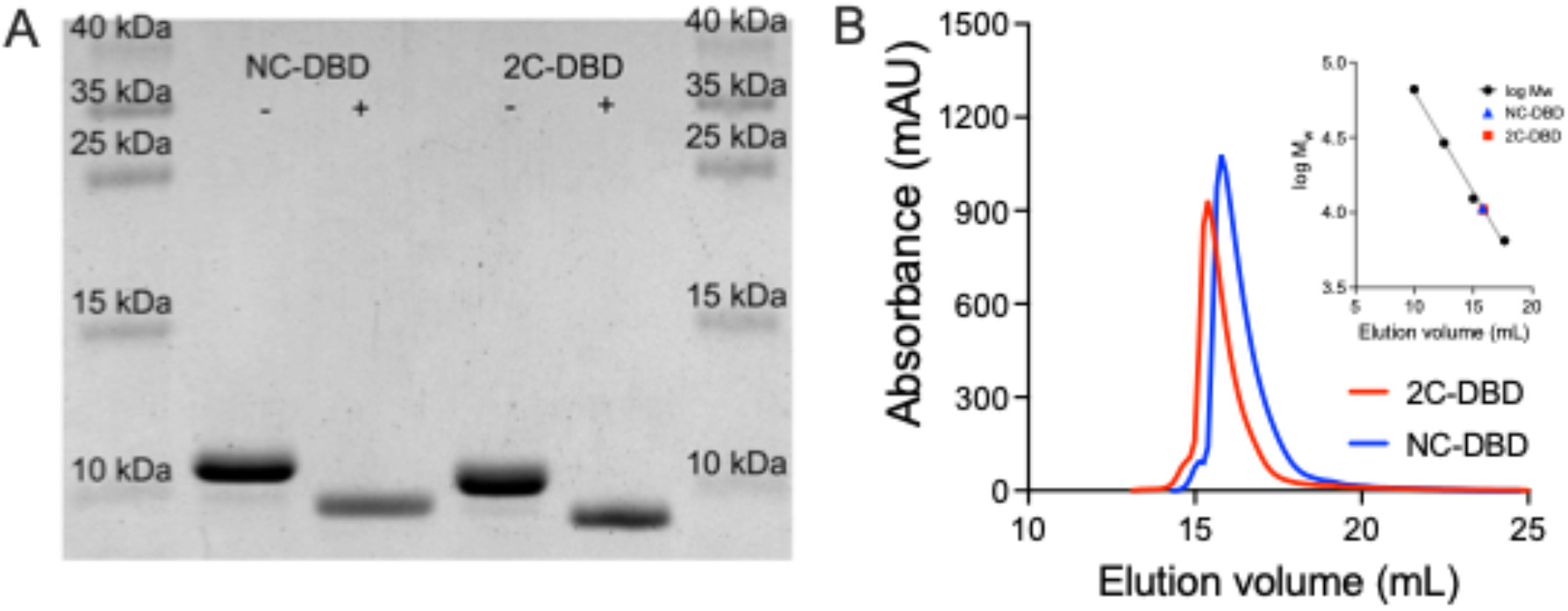
(A) SDS-PAGE of NC-DBD and 2C-DBD before (-) and after (+) His-tag cleavage. (B) Gel filtration chromatogram of NC-DBD and 2C-DBD. The molecular weight calibration curve is shown in the inset.

The oxidation state of as-purified 2C-DBD was determined using Ellman’s assay, which revealed that no free thiols are present (Table S1). This suggests that 2C-DBD either forms intramolecular or intermolecular disulfides. The oligomeric states of NC-DBD and 2C-DBD were determined by analytical gel filtration chromatography. Both NC-DBD and 2C-DBD had an elution volume that correspond to a 1.5-mer (Figure 3B). We interpret this result as meaning that both NC-DBD and 2C-DBD are monomers with some disordered regions that increase their hydrodynamic radius. The elution volume remained constant over a wide range of NC-DBD and 2C-DBD loading quantities (0.5 mg – 5 mg), indicating that the interaction between protomers in solution is weak and that dimerization does not occur at high protein concentration. To confirm gel filtration results and rule out weak protein-protein interactions, glutaraldehyde-based crosslinking was carried out. Glutaraldehyde is a nonspecific crosslinker that captures weak complexes [24]. As shown in Figure S3, the molecular weight of glutaraldehyde treated NC-DBD and 2C-DBD were identical to untreated samples, indicating that it is unlikely the domains dimerize in solution.

Analytical gel filtration data further indicates that reduction of the Cys residues by TCEP and DTT does not change the oligomeric state of 2C-DBD (Figure S4A). This suggests that the Cys pair forms an intramolecular disulfide when oxidized. The lack of an intermolecular disulfide was confirmed by SDS-PAGE. The migration distance of unreduced 2C-DBD was consistent with that of a monomer and identical to 2C-DBD that had been reduced (Figure S4B). Together, these data suggest that in *Gd*-NifA, the Cys residues in the IDL are redox active, however, they form an intramolecular disulfide and do not mediate DNA binding domain dimerization.

### Secondary structure analysis of NC-DBD and 2C-DBD

The secondary structure of both NC-DBD and 2C-DBD was investigated by circular dichroism (CD) spectroscopy (Figure 4). Both proteins are primarily α-helical, as expected based on the aforementioned structural predictions. Although the CD spectra of NC-DBD and 2C-DBD are similar, there are some significant differences. Analyzing the CD data using the CD fitting program K2D2 [25] indicates that the helical content in NC-DBD is 57%, whereas it is 27% for 2C-DBD. This is consistent with the prediction that the IDL region in 2C-DBD is disordered. The structure of 2C-DBD does not change between oxidizing and reducing conditions (Figure 4A). This indicates that formation of an intramolecular disulfide in 2C-DBD does not lead to significant structural changes in the IDL and that it is disordered in both reducing and oxidizing conditions. The melting point of NC-DBD and 2C-DBD was also determined, indicating that both proteins have nearly identical thermal stability (Figure 4B).

**Figure 4.**
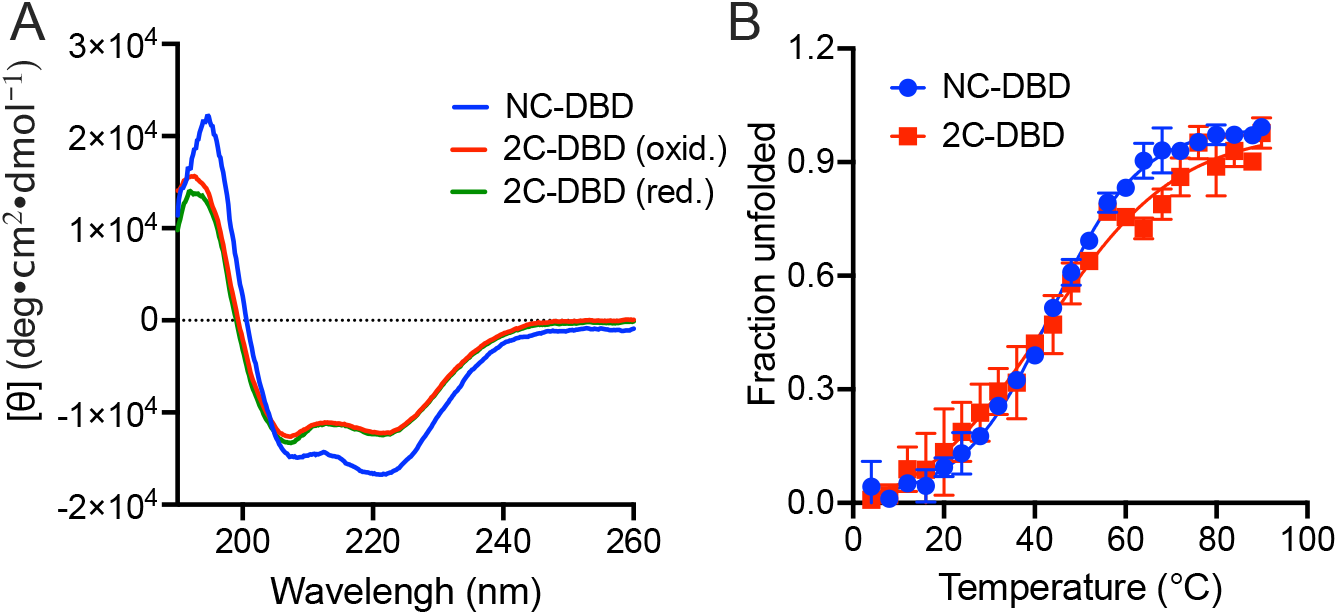
(A) CD spectra of NC-DBD and 2C-DBD. 2C-DBD was reduced with 2 mM TCEP. (B) Thermal denaturation curve of NC-DBD and 2C-DBD. The *T*_m_ for NC-DBD is 43.5 °C and that for 2C-DBD is 42.2 °C. The data represents an average of three independent measurements.

### DNA binding by the NifA DNA binding domain

To test whether the redox state of the Cys residues in the IDL influences DNA binding, we measured the binding affinity of NC-DBD and 2C-DBD with the UAS of *nifH* (5’-TGT-(N)_10_-ACA-3’) [26]. Duplexed nifH-UAS was labeled with a TAMRA fluorophore so that NifA binding could be measured by fluorescence anisotropy. Both NC-DBD and 2C-DBD bound to nifH-UAS in a dose dependent manner as evidenced by the increase in anisotropy as the protein concentration increased (Figure 5). Control experiments demonstrate that neither NC-DBD nor 2C-DBD bind to the fluorescent probe since addition of unlabeled nifH-UAS, which competes with fluorescently labeled nifH-UAS for DBD binding, causes a reversal of the anisotropy increase (Figure S5A). Furthermore, the nifH-UAS probe does not bind nonspecifically to non-DNA binding proteins such as BSA (Figure S5B).

**Figure 5.**
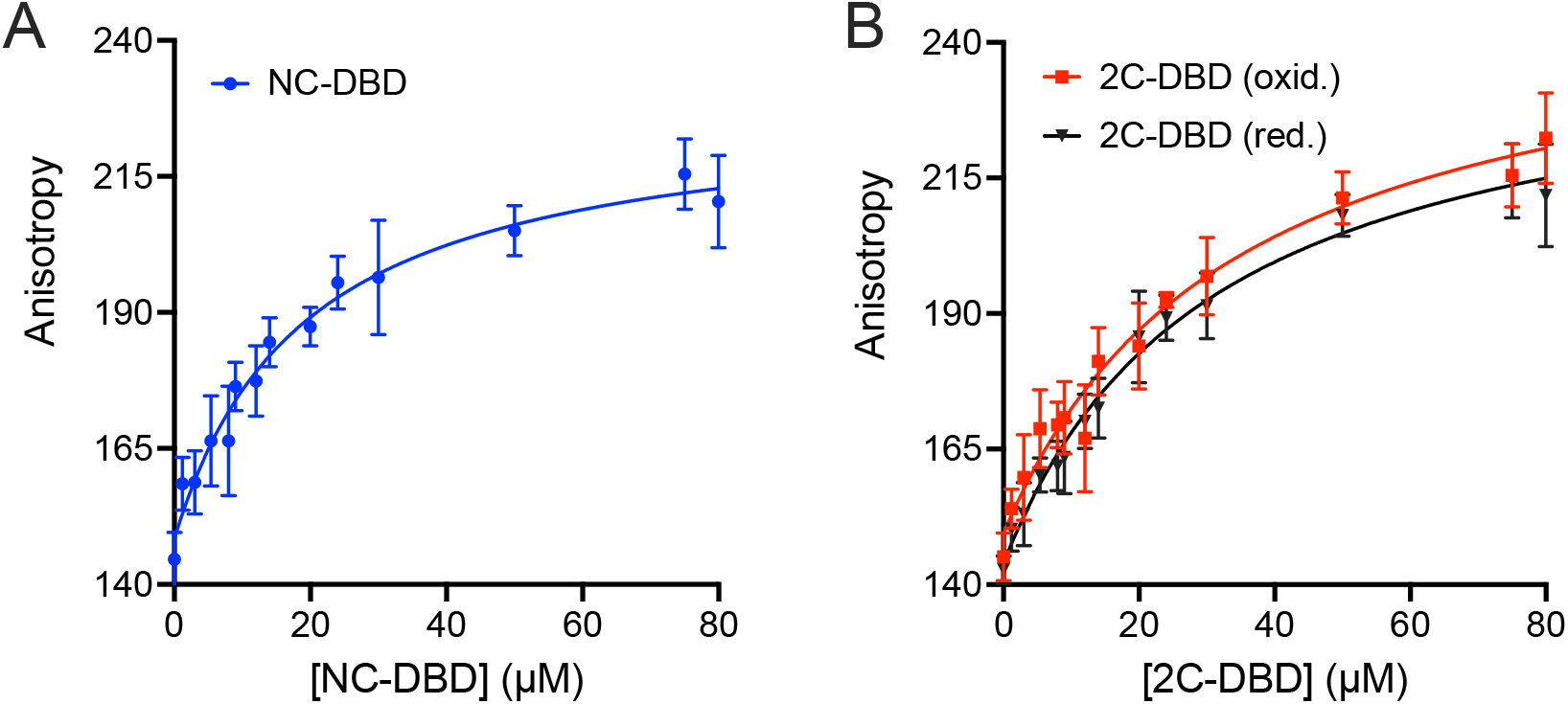
DNA binding domain binding to nifH-UAS for (A) NC-DBD and (B) 2C-DBD under oxidizing and reducing conditions. The *K*_d_ for NC-DBD binding is 20.0 ± 5.6 μM whereas that for 2C-DBD binding is 34.5 ± 8.4 μM and 31.5 ± 8.1 μM for oxidizing and reducing conditions, respectively.

NC-DBD and 2C-DBD binding to DNA follows a hyperbolic model. The binding affinity, *K*_d_, of NC-DBD was 20.0 ± 5.6 μM (Figure 5A). 2C-DBD bound to nifH-UAS with an affinity of 34.5 ± 8.4 μM. The binding affinity under reducing conditions (*K*_d_ = 31.5 ± 8.1 μM) is nearly unchanged, suggesting that breaking the intramolecular disulfide bond does not influence 2C-DBD’s structure in a way that affects DNA binding (Figure 5B). This result is consistent with the CD data that indicate that reduced and oxidized 2C-DBD are structurally indistinguishable and thus expected to bind DNA with the same affinity.

## Discussion

This work describes the purification and biophysical characterization of the DNA binding domain of NifA from the α-proteobacterium *G. diazotrophicus*. The domain’s secondary structure is mostly α-helical, which was expected based on its homology with DNA binding domains of other σ^54^ activators [8, 16, 27]. The IDL region immediately preceding the NifA DNA binding domain is disordered, making it similar to that of other σ^54^ activators such as NtrC1 [8], and NtrX [28], but different from the σ^54^ activators NtrC and NtrC4, which contain an additional helix, Helix A, prior to the HTH motif [8]. The *Gd*-NifA DNA binding domain is monomeric in solution. Interestingly, against our expectations, oxidation of the Cys pair did not promote dimer formation, which suggests that dimerization only occurs in presence of the palindromic DNA target. These results confirm previous reports that the DNA binding domain of σ^54^ activators that lack Helix A do not form a dimer in absence of DNA whereas those that have Helix A such as NtrC and NtrC4 form stable dimers [5, 8].

The binding affinities of NC-DBD and 2C-DBD towards the nifH-UAS are similar and not redox dependent. Reducing conditions do not increase the DNA binding affinity of 2C-DBD, indicating that redox sensing in NifA does not occur at the Cys pair upstream of the DNA binding domain. Surprisingly, the affinity of NC-DBD for DNA was higher than for 2C-DBD. We do not interpret this result as meaning that the IDL is a structural element that diminishes DNA binding. Instead, it is likely that the disordered IDL in 2C-DBD accesses conformations that interfere with DNA binding that it would not sample in the full-length protein. The binding affinity of NC-DBD and 2C-DBD is lower than that reported for NtrC and NtrC1 binding to their cognate UASs [16, 29] since both bind with low nM affinity. However, the *K*_d_ for DNA binding to *Gd*-NifA was similar to that reported for *Klebsiella pneumoniae* NifA binding to a nifH-UAS half site [30], which had a *K*_d_ of 200 μM. The reason for the difference in magnitude for DNA binding between NifA and NtrC/NtrC1 is unclear. It is possible that NifA inherently binds to upstream activator sequences less tightly than NtrC and NtrC1.

Redox-dependent activation of NifA in α-proteobacteria involves structural changes in response to reduction of either a metal cluster or a disulfide bond. Although our data does not provide evidence that the redox state of the Cys residues at the end of the IDL influence *Gd*-NifA activity, these residues may nonetheless be important for NifA function. Significant structural changes occur when σ^54^ activators undergo hexamerization, including large movements of the DNA binding domain relative to the AAA^+^ domain. It has been suggested that the IDL in σ^54^ activators is inherently flexible to support these structural changes [8]. Furthermore, the orientation of the DNA-binding domain relative to the AAA^+^ domain may influence binding to the RNA polymerase σ-factor [7, 31]. It is therefore possible that redox-dependent intramolecular disulfide bond breakage in the IDL regulates the flexibility of the IDL to facilitate the reorientation of the AAA^+^ domain in a redox dependent manner. Our group is currently investigating this hypothesis by characterizing full-length *Gd-*NifA.

## Abbreviations

CD: circular dichroism
DTT: Dithiothreitol
HEPES: 4-(2-hydroxyethyl)-1-piperazineethanesulfonic acid
IPTG: Isopropyl β-d-1-thiogalactopyranoside
MALDI-TOF: Matrix-assisted laser desorption/ionization – time of flight
SDS-PAGE: Sodium dodecyl sulfate–polyacrylamide gel electrophoresis
TCEP: Tris(2-carboxyethyl)phosphine
TRIS: Tris(hydroxymethyl)aminomethane
UAS: Upstream activator sequence

## Acknowledgements

The authors gratefully acknowledge Chris Kim, Andrew Lyon, and Marco Bisoffi for use of instrumentation.

## References

1. Dixon, R. & Kahn, D. (2004) Genetic regulation of biological nitrogen fixation, Nat Rev Microbiol. 2: 621–31.

2. Poza-Carrion, C., Jimenez-Vicente, E., Navarro-Rodriguez, M., Echavarri-Erasun, C. & Rubio, L. M. (2014) Kinetics of Nif gene expression in a nitrogen-fixing bacterium, J Bacteriol. 196: 595–603.

3. Fischer, H. M. (1994) Genetic regulation of nitrogen fixation in rhizobia, Microbiol Rev. 58: 352–86.

4. Demtroder, L., Pfander, Y., Schakermann, S., Bandow, J. E. & Masepohl, B. (2019) NifA is the master regulator of both nitrogenase systems in Rhodobacter capsulatus, Microbiologyopen. 8: e921.

5. Batchelor, J. D., Lee, P. S., Wang, A. C., Doucleff, M. & Wemmer, D. E. (2013) Structural mechanism of GAF-regulated sigma(54) activators from Aquifex aeolicus, J Mol Biol. 425: 156–70.

6. Rappas, M., Bose, D. & Zhang, X. (2007) Bacterial enhancer-binding proteins: unlocking sigma54-dependent gene transcription, Curr Opin Struct Biol. 17: 110–6.

7. Bush, M. & Dixon, R. (2012) The role of bacterial enhancer binding proteins as specialized activators of &#x03C3;54-dependent transcription, Microbiol Mol Biol Rev. 76: 497–529.

8. Vidangos, N., Maris, A. E., Young, A., Hong, E., Pelton, J. G., Batchelor, J. D. & Wemmer, D. E. (2013) Structure, function, and tethering of DNA-binding domains in sigma(5)(4) transcriptional activators, Biopolymers. 99: 1082–96.

9. Salazar, E., Diaz-Mejia, J. J., Moreno-Hagelsieb, G., Martinez-Batallar, G., Mora, Y., Mora, J. & Encarnacion, S. (2010) Characterization of the NifA-RpoN regulon in Rhizobium etli in free life and in symbiosis with Phaseolus vulgaris, Appl Environ Microbiol. 76: 4510–20.

10. Chen, S., Liu, L., Zhou, X., Elmerich, C. & Li, J.-L. (2005) Functional analysis of the GAF domain of NifA in Azospirillum brasilense: effects of Tyr&#x2192;Phe mutations on NifA and its interaction with GlnB, Mol Genet Genomics. 273: 415–422.

11. Oliveira, M. A., Aquino, B., Bonatto, A. C., Huergo, L. F., Chubatsu, L. S., Pedrosa, F. O., Souza, E. M., Dixon, R. & Monteiro, R. A. (2012) Interaction of GlnK with the GAF domain of Herbaspirillum seropedicae NifA mediates NH_4_^&#x002B;^-regulation, Biochimie. 94: 1041–7.

12. Martinez-Argudo, I., Little, R., Shearer, N., Johnson, P. & Dixon, R. (2004) The NifL-NifA System: a multidomain transcriptional regulatory complex that integrates environmental signals, J Bacteriol. 186: 601–10.

13. Zou, X., Zhu, Y., Pohlmann, E. L., Li, J., Zhang, Y. & Roberts, G. P. (2008) Identification and functional characterization of NifA variants that are independent of GlnB activation in the photosynthetic bacterium Rhodospirillum rubrum, Microbiology. 154: 2689–99.

14. Nishikawa, C. Y., Araujo, L. M., Kadowaki, M. A., Monteiro, R. A., Steffens, M. B., Pedrosa, F. O., Souza, E. M. & Chubatsu, L. S. (2012) Expression and characterization of an N-truncated form of the NifA protein of Azospirillum brasilense, Braz J Med Biol Res. 45: 113–7.

15. Oliveira, M. A., Baura, V. A., Aquino, B., Huergo, L. F., Kadowaki, M. A., Chubatsu, L. S., Souza, E. M., Dixon, R., Pedrosa, F. O., Wassem, R. & Monteiro, R. A. (2009) Role of conserved cysteine residues in Herbaspirillum seropedicae NifA activity, Res Microbiol. 160: 389–95.

16. Vidangos, N. K., Heideker, J., Lyubimov, A., Lamers, M., Huo, Y., Pelton, J. G., Ton, J., Gralla, J., Berger, J. & Wemmer, D. E. (2014) DNA recognition by a sigma(54) transcriptional activator from Aquifex aeolicus, J Mol Biol. 426: 3553–68.

17. Bertalan, M., Albano, R., de Pádua, V., Rouws, L., Rojas, C., Hemerly, A., Teixeira, K., Schwab, S., Araujo, J. & Oliveira, A. (2009) Complete genome sequence of the sugarcane nitrogen-fixing endophyte Gluconacetobacter diazotrophicus Pal5, BMC Genomics. 10: 450.

18. dos Santos, M. F., Muniz de Padua, V. L., de Matos Nogueira, E., Hemerly, A. S. & Domont, G. B. (2010) Proteome of Gluconacetobacter diazotrophicus co-cultivated with sugarcane plantlets, J Proteomics. 73: 917–31.

19. Owens, C. P. & Tezcan, F. A. (2018) Conformationally Gated Electron Transfer in Nitrogenase. Isolation, Purification, and Characterization of Nitrogenase From Gluconacetobacter diazotrophicus, Methods Enzymol. 599: 355–386.

20. Monteiro, R. A., Souza, E. M., Geoffrey Yates, M., Steffens, M. B., Pedrosa, F. O. & Chubatsu, L. S. (2003) Expression, purification, and functional analysis of the C-terminal domain of Herbaspirillum seropedicae NifA protein, Protein Expr Purif. 27: 313–8.

21. Medina, M. S., Bretzing, K. O., Aviles, R. A., Chong, K. M., Espinoza, A., Garcia, C. N. G., Katz, B. B., Kharwa, R. N., Hernandez, A., Lee, J. L., Lee, T. M., Lo Verde, C., Strul, M. W., Wong, E. Y. & Owens, C. P. (2021) CowN sustains nitrogenase turnover in the presence of the inhibitor carbon monoxide, J Biol Chem. 296: 100501.

22. Baek, M., DiMaio, F., Anishchenko, I., Dauparas, J., Ovchinnikov, S., Lee, G. R., Wang, J., Cong, Q., Kinch, L. N., Schaeffer, R. D., Millan, C., Park, H., Adams, C., Glassman, C. R., DeGiovanni, A., Pereira, J. H., Rodrigues, A. V., van Dijk, A. A., Ebrecht, A. C., Opperman, D. J., Sagmeister, T., Buhlheller, C., Pavkov-Keller, T., Rathinaswamy, M. K., Dalwadi, U., Yip, C. K., Burke, J. E., Garcia, K. C., Grishin, N. V., Adams, P. D., Read, R. J. & Baker, D. (2021) Accurate prediction of protein structures and interactions using a three-track neural network, Science. 373: 871–876.

23. Barozet, A., Molloy, K., Vaisset, M., Zanon, C., Fauret, P., Simeon, T. & Cortes, J. (2021) MoMA-LoopSampler: A web server to exhaustively sample protein loop conformations, Bioinformatics.

24. Migneault, I., Dartiguenave, C., Bertrand, M. J. & Waldron, K. C. (2004) Glutaraldehyde: behavior in aqueous solution, reaction with proteins, and application to enzyme crosslinking, Biotechniques. 37: 790–6, 798-802.

25. Perez-Iratxeta, C. & Andrade-Navarro, M. A. (2008) K2D2: estimation of protein secondary structure from circular dichroism spectra, BMC Struct Biol. 8: 25.

26. Morett, E. & Buck, M. (1988) NifA-dependent in vivo protection demonstrates that the upstream activator sequence of nif promoters is a protein binding site, Proc Natl Acad Sci U S A. 85: 9401–5.

27. Batchelor, J. D., Doucleff, M., Lee, C. J., Matsubara, K., De Carlo, S., Heideker, J., Lamers, M. H., Pelton, J. G. & Wemmer, D. E. (2008) Structure and regulatory mechanism of Aquifex aeolicus NtrC4: variability and evolution in bacterial transcriptional regulation, J Mol Biol. 384: 1058–75.

28. Fernandez, I., Cornaciu, I., Carrica, M. D., Uchikawa, E., Hoffmann, G., Sieira, R., Marquez, J. A. & Goldbaum, F. A. (2017) Three-Dimensional Structure of Full-Length NtrX, an Unusual Member of the NtrC Family of Response Regulators, J Mol Biol. 429: 1192–1212.

29. Sevenich, F. W., Langowski, J., Weiss, V. & Rippe, K. (1998) DNA binding and oligomerization of NtrC studied by fluorescence anisotropy and fluorescence correlation spectroscopy, Nucleic Acids Res. 26: 1373–81.

30. Ray, P., Smith, K. J., Parslow, R. A., Dixon, R. & Hyde, E. I. (2002) Secondary structure and DNA binding by the C-terminal domain of the transcriptional activator NifA from Klebsiella pneumoniae, Nucleic Acids Res. 30: 3972–80.

31. De Carlo, S., Chen, B., Hoover, T. R., Kondrashkina, E., Nogales, E. & Nixon, B. T. (2006) The structural basis for regulated assembly and function of the transcriptional activator NtrC, Genes Dev. 20: 1485–95.

