## Supporting information for "Purification and biochemical characterization of the DNA binding domain of the nitrogenase transcriptional activator NifA from *Gluconacetobacter diazotrophicus*"

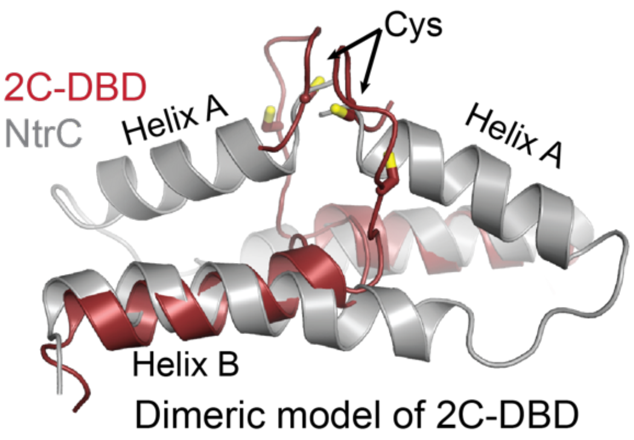


**Figure S1.** Model of dimeric 2C-DBD aligned with dimeric NtrC (pdb id: 1ntc). Possible orientations of the flexible IDL region in 2C-DBD were modeled using the MoMA loop sampler, revealing IDL orientations that place Cys resides of different protomers in proximity to each other. The 2C-DBD dimer was created by aligning two copies of 2C-DBD with the two chains in the NtrC dimer. For clarity, only the IDL, Helix A (present only in NtrC), and Helix B are depicted.

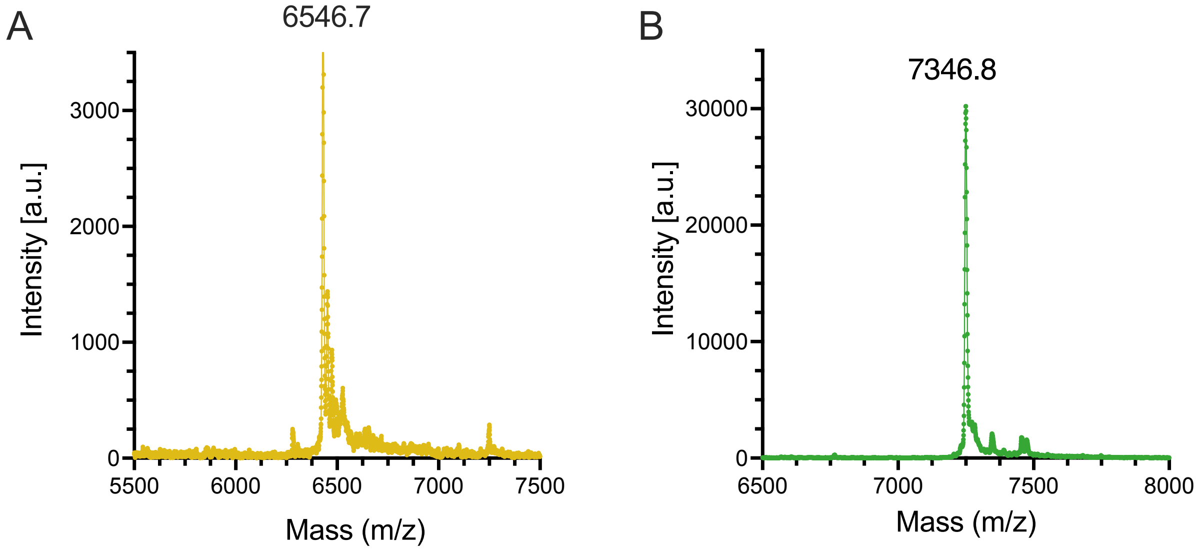


**Figure S2.** MALDI-TOF of (A) NC-DBD and (B) 2C-DBD after His-tag cleavage. The expected masses are 6427.4 g/mol and 7247.3 g/mol for NC-DBD and 2C-DBD, respectively.

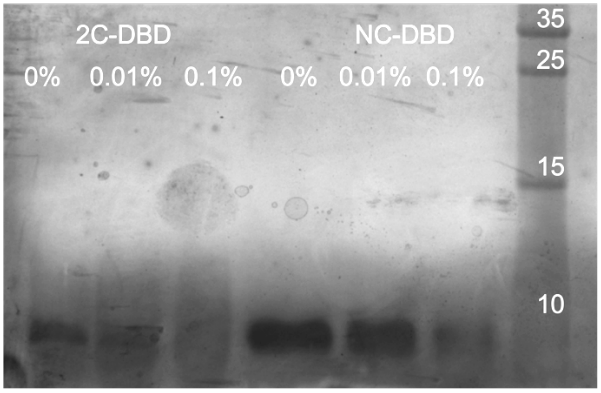


**Figure S3.** Glutaraldehyde crosslinking of 2C-DBD and NC-DBD indicating that no dimer or higher order oligomers are formed in presence of the nonspecific crosslinker glutaraldehyde. Glutaraldehyde concentrations are listed above each lane. The gel was silver stained to detect potential small amounts of dimer which would appear at 13 kDa for NC-DBD and 14.5 kDa for 2C-DBD. The gel is representative of three independent replicates. The rightmost lane represents the molecular weight marker with standards in kDa.

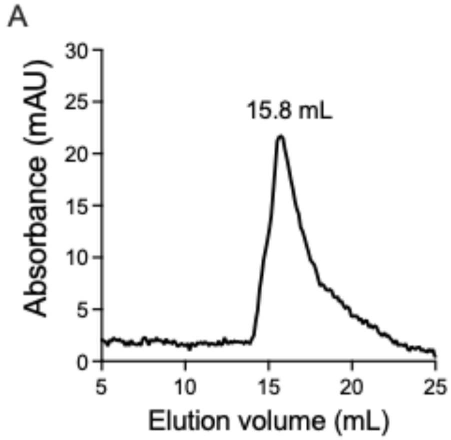


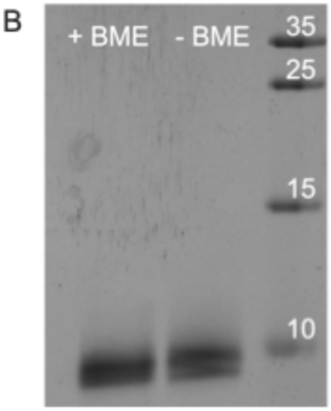


**Figure S4.** (A) Analytical gel filtration chromatograms of 2C-DBD run in presence of 5 mM TCEP. Results are identical when 10 mM DTT was used as a reducing agent instead of TCEP. (B) SDS-PAGE of 2C-DBD that was reduced with 5% BME and 2C-DBD that was not reduced. The migration distance is the same for both samples, indicating that the Cys residues do not form an intermolecular disulfide. The rightmost lane is the molecular weight marker with standards in kDa.

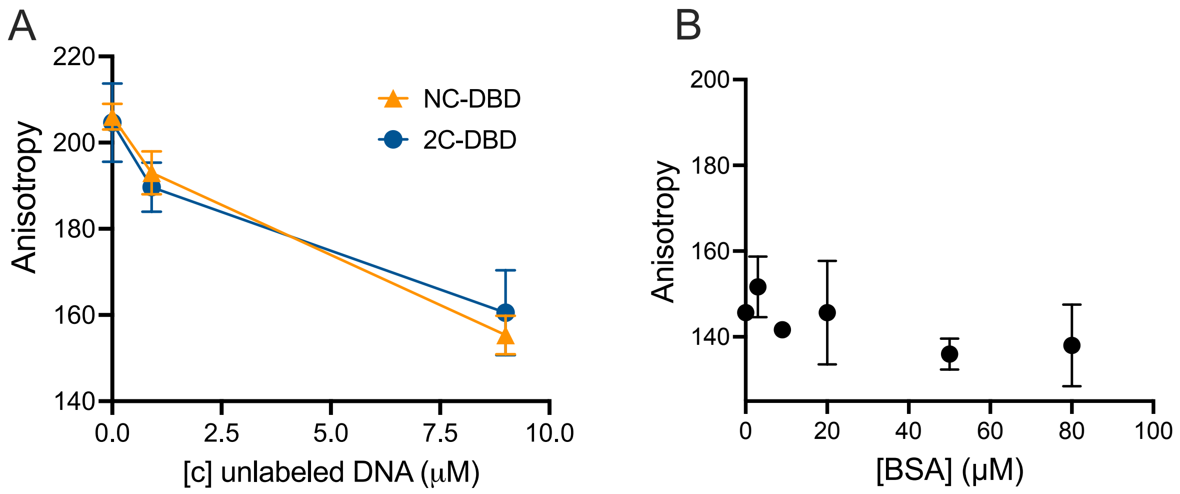


**Figure S5.** (A) Controls indicating that the DNA binding domain does not bind nonspecifically to the fluorescent probe. The protein concentration in both experiments was 50 μM and the labeled DNA concentration was 900 nM. (B) Control experiments demonstrating that the nifH-UAS probe does not bind nonspecifically to BSA.

**Table S1.** Ellman's assay demonstrating that 2C-DBD does not have free Cys in its as-purified state. NC-DBD served as a negative control as it does not have any Cys residues in its amino acid sequence. BSA severed as a positive control as it is expected to have a single free Cys residue. Data represents averages of duplicate measurements.

| **Protein** | **Concentration (μM)** | **Free Cys concentration (μM)** |
|---|---|---|
| 2C-DBD | 150 | 4.7 |
| 2C-DBD | 50 | 4.5 |
| NC-DBD | 150 | 3.7 |
| NC-DBD | 50 | 4.8 |
| BSA | 200 | 206.4 |
| BSA | 50 | 44.6 |
